# Single cell transcriptomics reveal disrupted kidney filter cell-cell interactions after early and selective podocyte injury

**DOI:** 10.1101/2020.07.30.229666

**Authors:** Abbe R. Clark, Jamie Marshall, Yiming Zhou, Monica S. Montesinos, Haiqi Chen, Lan Nguyen, Fei Chen, Anna Greka

## Abstract

The health of the kidney filtration barrier requires communication between podocytes, endothelial cells and mesangial cells. Disruption of these cell-cell interactions is thought to contribute to disease progression in chronic kidney diseases (CKD). We recently demonstrated that podocyte ablation via doxycycline-inducible deletion of an essential endogenous molecule, CTCF (iCTCF^pod-/-^), is sufficient to drive progressive CKD. However, the earliest events connecting podocyte injury to disrupted intercellular communication within the kidney filter remain unclear. Here we performed single-cell RNA sequencing of kidney tissue from iCTCF^pod-/-^ mice after one week of doxycycline induction to generate a map of the earliest transcriptional effects of podocyte injury on cell-cell interactions at single cell resolution. A subset of podocytes showed the earliest signs of injury due to disrupted gene programs for cytoskeletal regulation and mitochondrial function. Surviving podocytes upregulated Col4a5, causing reactive changes in integrin expression in endothelial populations and mesangial cells. Intercellular interaction analysis revealed several receptor-ligand-target gene programs as drivers of endothelial cell injury and abnormal matrix deposition. This analysis reveals the earliest disruptive changes within the kidney filter, pointing to new, actionable targets within a therapeutic window that may allow us to maximize the success of much needed therapeutic interventions for CKD.

## Introduction

Chronic kidney diseases (CKD) affect 700 million people globally and yet specific therapies are lacking (1). Many kidney diseases originate in the glomerulus, the filtration unit of the kidney. The glomerulus consists of (i) podocytes, specialized postmitotic cells with elaborate foot processes that interdigitate forming slit diaphragms and wrapping around glomerular capillaries, (ii) endothelial cells that lie opposite podocytes on a shared glomerular basement membrane (GBM), (iii) mesangial cells that form a matrix providing structural support for the glomerulus and (iv) parietal epithelial cells (PEC) that line Bowman’s Capsule (2). Podocyte injury in particular leads to many highly prevalent, progressive kidney diseases, including diabetic kidney disease (DKD), focal segmental glomerulosclerosis (FSGS) and nephrotic syndrome (both idiopathic and genetic). The canonical pattern of injury results in the loss of interdigitating podocyte foot processes, known as foot process effacement, caused by a rearrangement of the actin cytoskeleton. This leads to a disruption of the slit diaphragm, the physical barrier that functions as a filter, followed by podocyte detachment or death (3, 4). On the other hand, in addition to intact podocytes, the formation and maintenance of the glomerular filtration barrier requires intraglomerular communication, tightly controlled by a series of autocrine and paracrine signaling mechanisms. For example, VEGFA is a pro-survival signal for endothelial cells secreted by podocytes and PDGFB is a pro-survival signal for mesangial cells secreted by endothelial cells (5). Disruption of these cell-cell interactions are frequently observed in a host of glomerular diseases, including FSGS and DKD (6). Therefore, identifying the earliest disruptive changes to the glomerulus may offer novel targets, and the opportunity to optimize therapeutic success for the treatment of kidney diseases.

Single-cell RNA sequencing (scRNAseq) has revolutionized our ability to study individual cell types of complex tissues and cell states after specific perturbations (7, 8). Recent scRNAseq studies in kidney have provided insight into the transcriptional profiles of kidney cell types in healthy mice and human samples, as well as in some disease states, including DKD and lupus nephritis (9–12). However, detailed studies of glomerular cell states have been limited by the small number of cells available for analysis. Furthermore, the earliest cell type-specific changes that occur in all glomerular cells upon podocyte injury have not been monitored and are yet to be defined.

We recently showed that podocyte ablation via podocyte-specific inducible deletion of an essential endogenous protein, CTCF, leads to progressive proteinuric kidney disease and CKD (13). Historically, CKD mouse models were generated by inducing kidney injury by exogenous toxins or surgical interventions, suboptimal systems that fall short of recapitulating sequential mechanistic changes. Deletion of CTCF, an essential endogenous molecule, led to rapid podocyte loss, severe progressive albuminuria, bone mineral metabolism changes, kidney failure and premature death (13). The iCTCF^pod-/-^ model provides the unique opportunity to study changes in intraglomerular cell-cell interactions as a consequence of induced podocyte injury. As previously shown, podocyte CTCF expression was undetectable at the earliest timepoint, at one week of doxycycline-mediated Cre induction in iCTCF^pod-/-^ mice (13). Significant and progressive podocyte loss, as measured by histologic analysis, started at two weeks after Cre induction. For the current study, we chose to perform a detailed analysis at the one week time point to identify the earliest changes in intraglomerular cell-cell interactions driven by podocyte injury that ultimately lead to CKD.

## Results

### Single-cell profiling of over 29,000 glomerular-enriched kidney cells

Current single cell protocols for whole kidney identify < 2.5% glomerular cells (9) and while a purified glomerular preparation using magnetic beads enriches for this population (14), it fails to capture other cell types of the kidney that may be of interest. To extend these findings and develop a detailed understanding of cell-cell interactions within the glomerulus in the context of the entire cellular landscape of the kidney, before and after podocyte injury, we used a sieving method to simultaneously enrich glomeruli and capture additional kidney cell types.

To identify the early transcriptional effects of podocyte ablation in a cell type-specific manner, scRNAseq was performed with kidney tissue from WT and iCTCF^pod-/-^ mice collected by serial sieving after one week of doxycycline treatment (Figure 1A). Kidney tissue from four WT animals (four biological replicates) and four iCTCF^pod-/-^ animals yielded a total of 14783 WT and 14727 iCTCF^pod-/-^ cells profiled after filtering (Supplemental Figure 1A, 1B; Supplemental Tables 1, 2). Data was normalized to remove effects due to the number of unique molecular identifiers (UMI) and percent mitochondrial reads. After integrating WT and iCTCF^pod-/-^ samples, we used a low resolution of clustering to detect nine clusters (Figure 1B). Biological replicates of WT and iCTCF^pod-/-^ samples were distributed amongst all clusters (Supplemental Figure 2A-C). All cell types of the glomerulus, as well as additional kidney cell types, were identified using established and data-derived markers (Figure 1C; Supplemental Table 3; Supplemental Figure 3A).

**Figure 1.**
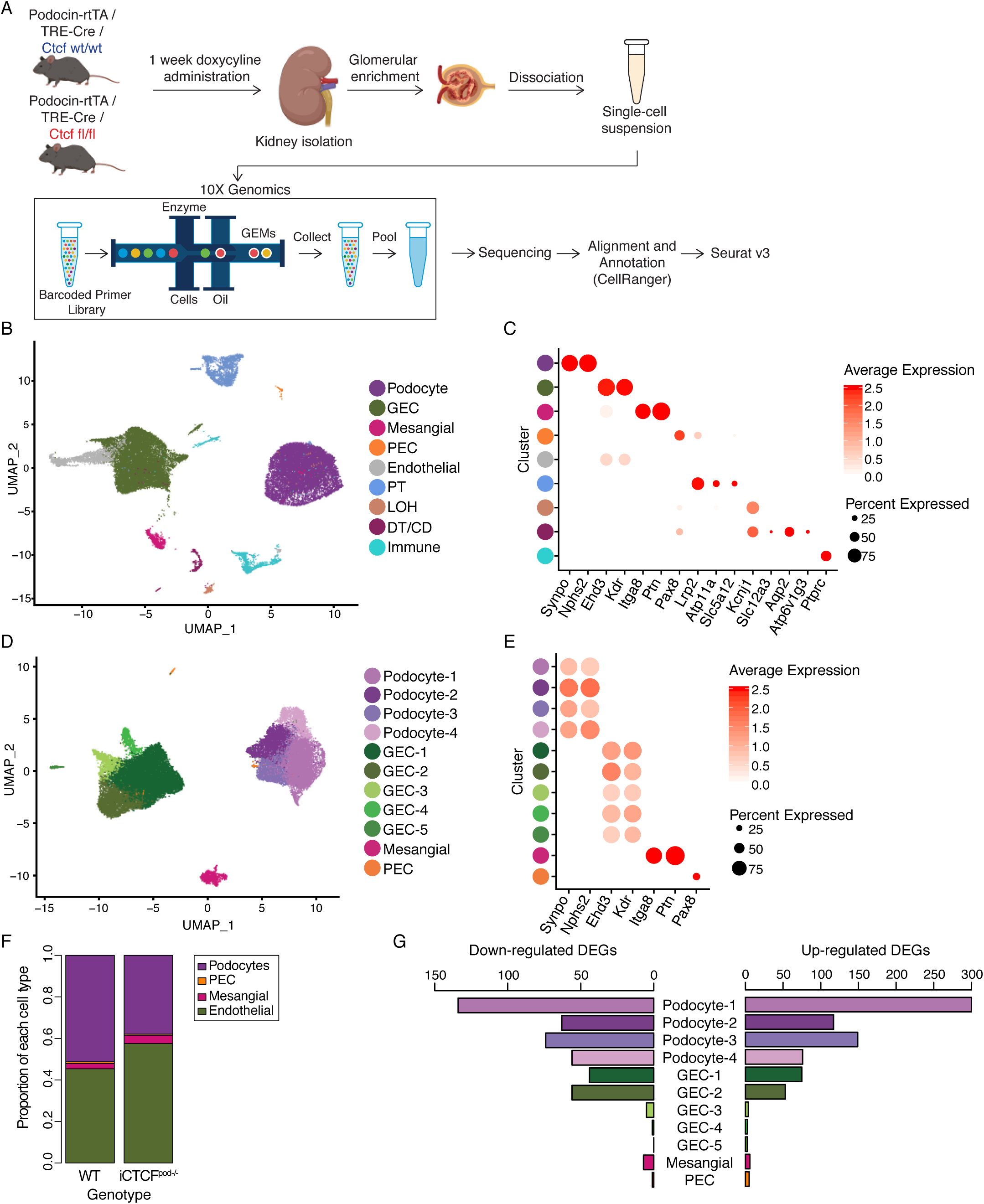
Single-cell RNA sequencing of WT and iCTCF^pod-/-^ kidneys shows that endothelial cells are most affected by podocyte injury. (A) Experimental design. WT and iCTCF^pod-/-^ mice were treated with doxycycline for 1 week to induce deletion of *Ctcf* specifically in podocytes. After 1 week, kidneys were isolated, and glomeruli were enriched using serial sieving. Enzymatic digestion was used to obtain a single-cell suspension that was subsequently sequenced on the 10X Genomics platform. Downstream data analysis was performed using CellRanger and Seurat. (B) UMAP plot of all cells isolated from WT and iCTCF^pod-/-^ glomerular-enriched kidney fractions. (C) Expression of cell-type specific markers used for cluster identification in A presented as a dot plot. Color represents the average expression level of the gene in the specified cluster. WT and iCTCF^pod-/-^ cells split into 10 clusters and all major cell types of the kidney were identified. (D) UMAP plot of cells of glomerular origin that were isolated and reclustered. (E) Expression of cell-type specific markers used for cluster identification in C. WT and iCTCF^pod-/-^ glomerular cells split into 11 clusters. (F) Proportion of WT or iCTCF^pod-/-^ cells per glomerular cell type. (G) Barplot of the number of differentially expressed genes (DEGs) in each cluster comparing iCTCF^pod-/-^ vs. WT cells. Genes were defined as differentially expressed if they were expressed in at least 10% of cells, had a log fold-change of 0.01 and an adjusted p-value of < 0.05.

Glomerular cells, representing a total 82.1% of all recovered cells (36.9% podocytes, 42.1% glomerular endothelial cells (GECs), 2.7% mesangial cells, 0.36% parietal epithelial cells (PECs)) were isolated and re-clustered (Figure 1D). Four clusters of podocytes, five clusters of GECs and one cluster each of mesangial cells and PECs, expressing canonical cell-type markers, were identified (Figure 1E; Supplemental Figure 3B). Biological replicates of WT and iCTCF^pod-/-^ samples were distributed throughout all clusters (Supplemental Figure 2D-F; Supplemental Table 4). As anticipated, given that podocyte-specific CTCF deletion leads to histologically detectable podocyte loss at 2 weeks (13), iCTCF^pod-/-^ samples contained a lower percentage of podocytes than WT samples (Figure 1F; Supplemental Table 5). iCTCF^pod-/-^ samples had 13.1% fewer podocytes, 11.9% more GEC and 1.5% more mesangial cells than WT samples (Supplemental Table 5). PEC contributed < 1% to either of the iCTCF^pod-/-^ or WT samples. This suggests that disruption of transcriptional programs critical for podocyte survival precedes histologically detectable podocyte injury and loss and highlights the power of scRNAseq in discerning subtle changes that can be missed by histology.

### Podocyte-specific inducible CTCF deletion leads to gene expression changes in all glomerular cell types

Differential expression analysis was performed in each of 11 clusters comparing iCTCF^pod-/-^ vs. WT cells (Supplemental Table 6). Genes were considered differentially expressed if found in at least 10% of cells in a given cluster, with a minimum absolute log fold-change of 0.1 and an adjusted p-value < 0.05. *Ctcf* was differentially expressed in each of the four podocyte clusters, with average log fold changes of -0.390, -0.189, - 0.331, -0.303, respectively. To compare the number of differentially expressed genes among the 11 clusters, we downsampled the numbers of cells in each cluster and repeated the differential expression analysis. The podocyte clusters had the most differentially expressed genes, followed by GEC-1and GEC-2 (Figure 1G). The remaining three clusters of GECs, along with the mesangial cells and PECs, had far fewer differentially expressed genes (Figure 1G), suggesting that GEC-1 and GEC-2, among all glomerular clusters, were most affected by the sequelae of CTCF-deletion-driven podocyte injury.

### Disease-associated gene programs identified in specific podocyte and endothelial cell clusters

We first sought to examine how the individual podocyte clusters respond to injury. A Venn diagram of the differentially expressed genes in each of the four podocyte clusters revealed that Podocyte-1 had the most uniquely differentially expressed genes of the four podocyte clusters (Figure 2A; Supplemental Table 7). This observation that one out of four podocyte clusters was more prominently affected, is in agreement with prior histologic data showing that not all podocytes are affected with equal severity in the face of injury and highlights the power of scRNAseq to molecularly characterize the heterogeneity of cell states within the same cell type (15).

**Figure 2.**
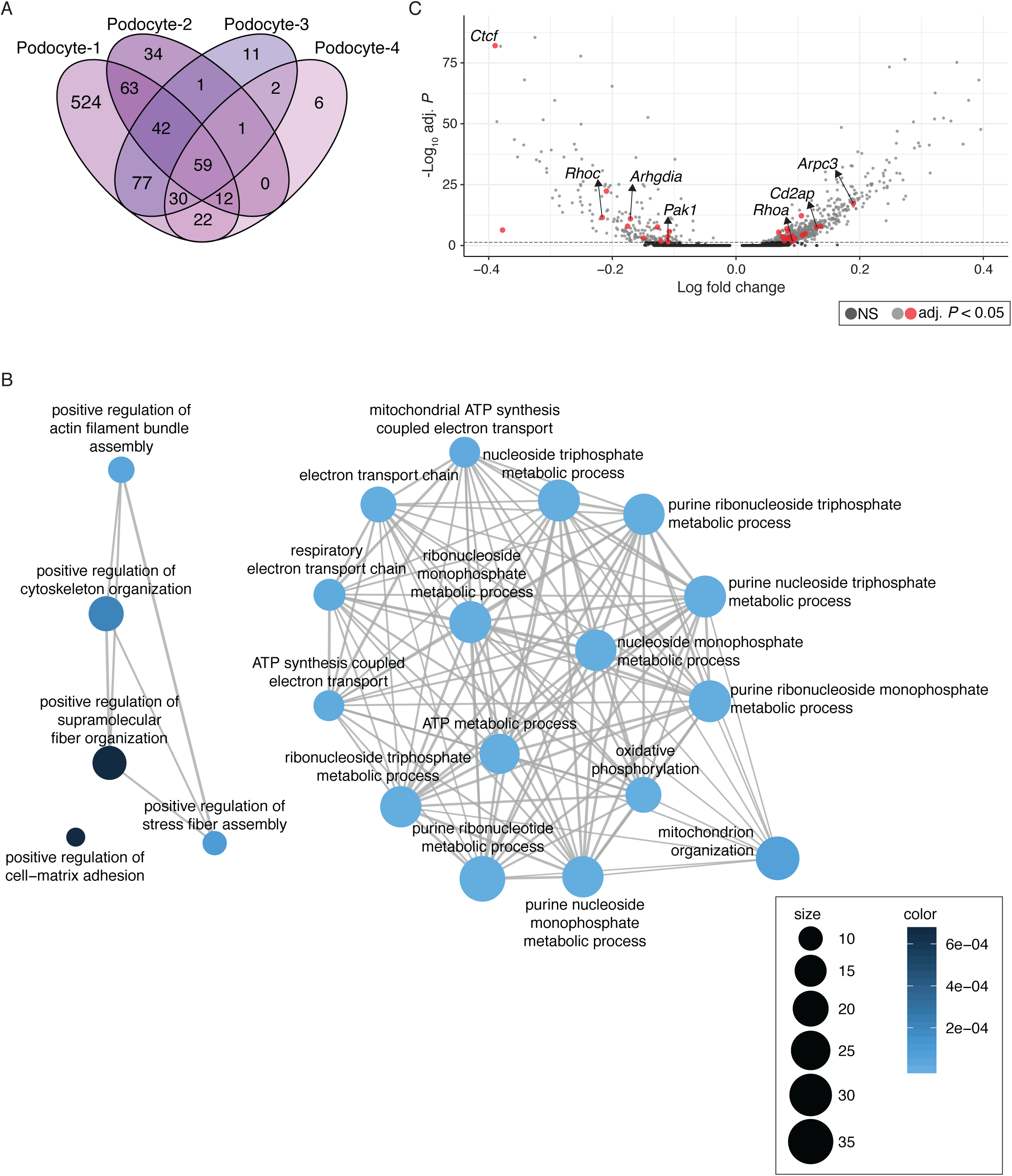
Gene programs of focal adhesions, cytoskeleton organization and mitochondrial functions enriched in podocytes. (A) Venn diagram of the differentially expressed genes identified in each of the four podocyte sub-clusters. (B) Visualization of the over-representation analysis of genes uniquely differentially expressed in Podocyte-1 presented as a network, or enrichment map. Overlapping gene sets cluster together. (C) Volcano plot of differentially expressed genes (comparing iCTCF^pod-/-^ cells v. WT cells) in Podocyte-1. All genes from adhesion and cytoskeleton organization gene programs are colored in red, with genes of interest labeled. Significantly differentially expressed genes are represented by light grey and red dots. Genes in dark grey are not significantly differentially expressed. The y-axis has been limited to 100.

To identify gene programs enriched as a consequence of CTCF loss in the podocyte-1 cluster, we performed an over-representation analysis (Supplemental Table 8). We visualized the top enriched terms with an enrichment map to cluster mutually overlapping gene sets (Figure 2B). A prominent group of enriched terms was mitochondrial functions including ATP synthesis, mitochondrial organization, electron transport chain and oxidative phosphorylation (Figure 2B). These data extend recent work pointing to mitochondrial dysfunction as a sign of podocyte injury (16). In addition, human genetics have pointed to the importance of mitochondrial functions in podocytes including several mutations in the CoQ biosynthesis pathway (PDSS1, PDSS2, COQ2, COQ6, ADCK4) that cause nephrotic syndrome, mainly in children (17, 18). It has therefore been postulated that podocyte mitochondrial dysfunction may represent a prominent cell state associated with all diseases that stem from podocyte loss (16). Our data provide support for this notion at single cell resolution, suggesting that mitochondrial dysfunction may represent the earliest injury state in a specific population of podocytes leading to podocyte loss.

One of the main groups of enriched terms from the overrepresentation analysis (Figure 2B) was cytoskeletal organization as well as a related single enriched term for cell-matrix adhesion. The actin cytoskeleton plays an essential role in maintaining the podocytes’ unique and complex structure and adhesion to the GBM is essential for podocyte function (19). Mature focal adhesions contain hundreds of proteins that link the actin cytoskeleton, receptor-matrix binding, intracellular signal transduction and actin polymerization. One of the most downregulated genes in Podocyte-1 was *Rhpn1* (Supplemental Figure 4A), an essential component for establishing podocyte cytoskeleton dynamics and maintaining podocyte foot process architecture (20). The Arp2/3 complex component *Arpc3*, a driver of actin polymerization, was upregulated in the Podocyte-1 cluster (Figure 2C; Supplemental Figure 4B). A gene for an additional component of the complex, *Actr2* was also upregulated in both Podocyte-1 and Podocyte-4. In addition, *Wasl*, whose protein product, N-WASP, activates the Arp2/3 complex and is required for the maintenance of podocyte foot processes *in vivo* (21), was also upregulated in Podocyte-1, Podocyte-3 and Podocyte-4. The cytoskeletal regulator *Arhgdia*, whose deletion is associated with nephrotic syndrome in mice (22), was downregulated in Podocyte-1 (Figure 2C). The expression levels of two Rho GTPases that are well established regulators of the actin cytoskeleton and cell adhesion dynamics (23) were also altered: *RhoA* was upregulated and *RhoC* was downregulated, as was a central downstream effector of Rac1, *Pak1*, consistent with prior studies (15). Further, Cd2ap, a critical podocyte actin cytoskeleton component (19, 24) was upregulated in Podocyte-1. Together, these changes suggested that one of the earliest podocyte responses to CTCF deletion-driven injury is to alter critical components of the actin cytoskeleton in a struggle to maintain attachment to the GBM and thus survive the injury. In addition, scRNAseq identified specific mediators of this response matched to a specific population of podocytes, pointing to putative targets for early therapeutic intervention.

The GBM is a meshwork of extracellular matrix proteins, situated between podocytes and GECs, that provides structural support for the glomerular capillaries, harbors ligands for receptors on the surface of the adjacent GECs, podocytes and mesangial cells and contributes to glomerular filter selectivity (25). One of the critical components of the GBM is collagen type IV □5 (Col4a5); mutations in this gene cause Alport syndrome and FSGS in humans (25, 26). Variants in *Col4a3*, another structural component of the GBM, were recently identified by a comprehensive genome wide association study (GWAS) in diabetic kidney disease (DKD) (27). Therefore, changes to GBM components as a consequence of podocyte injury are broadly relevant to many kidney diseases. We asked if expression of genes encoding structural components of the GBM were disrupted in our model. *Col4a5* was upregulated in iCTCF^pod-/-^ podocytes (with a log fold change of 0.130; Supplemental Figure 4C; Supplemental Table 6). Ligand-receptor analysis suggested that *Col4a5* upregulation in Podocyte-1 leads to increased interactions with cells of all five GEC clusters and mesangial cells through several integrins (Figure 3A). We validated the observed *Col4a5* upregulation, with spatial resolution, by Hybridization Chain Reaction (HCR), a method that generates single-molecule fluorescence via *in situ* hybridization (28) (Figure 3B; Supplemental Figure 5). The expression of *Col4a5* in *Nphs2-*expressing podocytes was significantly increased (p < 0.0001; Welch-corrected two-tailed t-test) (Figure 3C). This result confirmed that individual gene data derived from the single cell transcriptomic experiment could be independently validated, with spatial resolution, bolstering the validity of the conclusions drawn by computational analyses. Furthermore, these data highlight that the key response to injury by a specific population of podocytes is to upregulate collagen production, which may account for the prominent or thick GBM observed in progressive diseases such as DKD (27).

**Figure 3.**
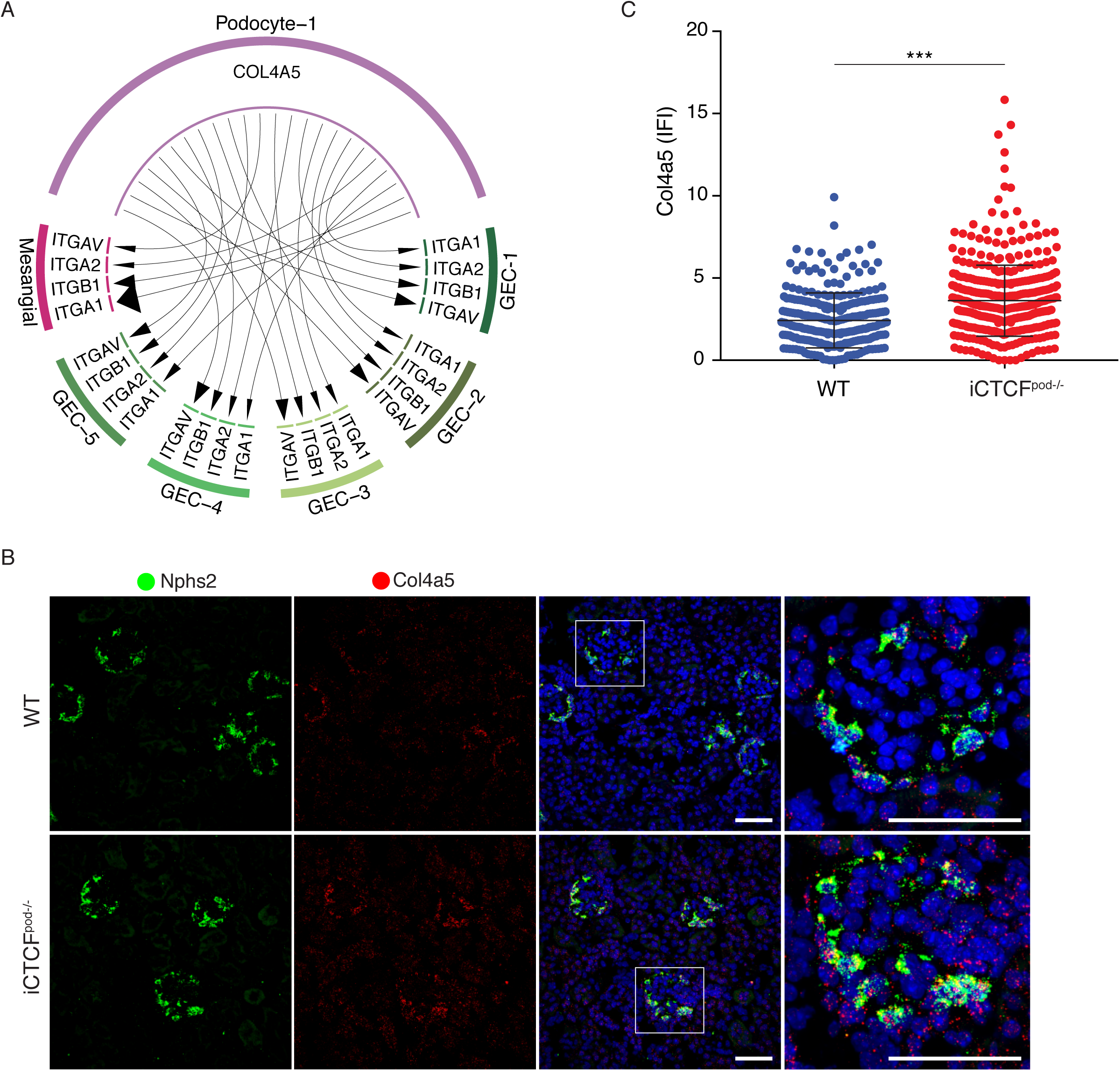
Podocyte cluster upregulates *Col4a5*. (A) *Col4a5* was significantly upregulated in iCTCF^pod-/-^ Podocyte-1 cells compared to WT Podocyte-1 cells. COL4A5 receptors were strongly expressed in mesangial cells and GEC subclusters 1-4. Size of the arrow head represents the relative expression of the receptor. (B) *Col4a5* expression in glomeruli of WT and iCTCF^pod-/-^ animals as evaluated by hybridization chain reaction (HCR). (C) Quantification of HCR. *Col4a5* was significantly upregulated in iCTCF^pod-/-^ *Nphs2*-expressing cells compared to WT *Nphs2*-expressing cells (p-value <0.0001; two-tailed T-test with Welch’s correction). Four WT and four iCTCF^pod-/-^ mice were evaluated. All glomeruli from three 40X images were quantified. Scale bars are 50 μm.

Turning our attention to other glomerular cells, we found that GEC-1 and GEC-2 had nearly as many differentially expressed genes as podocyte clusters 2-4, whereas the remaining GEC clusters, mesangial cells and PEC had fewer differentially expressed genes. We concluded that these two endothelial clusters were most affected by podocyte injury. GEC-1 expressed over 600 uniquely differentially expressed genes (Supplemental Figure 6A; Supplemental Table 9). Over-representation analysis revealed that the uniquely differentially expressed genes in GEC-1 were enriched in gene programs for cell migration and adhesion (Supplemental Figure 6B, C; Supplemental Table 10). These analyses revealed the earliest transcriptional changes that occur in GECs as a response to podocyte injury.

### Modelling intercellular communication reveals key interactions between cell types in response to early podocyte injury

Having identified disrupted gene programs in several distinct cell clusters, we next sought to understand how glomerular cell-cell crosstalk was influenced by podocyte injury. We first probed the list of differentially expressed genes in podocyte, GEC and mesangial cell clusters. We found that the expression of two key autocrine pro-survival ligands, *Vegfa* and *Pdgfb*, as well as the expression of the PDGFB receptor, *Pdgfrb* were disrupted (Supplemental Table 6). *Vegfa* expression was decreased in all podocyte clusters (average log-fold change between -0.034 and -0.149). In contrast, *Pdgfb* expression was increased in all GEC clusters (average log-fold change between 0.031 and 0.140) and the receptor, *Pdgfrb* was upregulated in mesangial cells (average log-fold change of 0.155). This analysis suggested that podocyte injury leads to decreased expression of the pro-survival ligand *Vegfa*, negatively affecting GECs. To compensate, GECs may upregulate the pro-survival ligand *Pdgfb*, triggering mesangial cells to upregulate the receptor *Pdgfrb*.

To more deeply investigate ligand-receptor interactions and their putative target genes, we used NicheNet, a novel algorithm that infers how ligand-receptor interactions derived from expression data may affect specific targets by integrating pre-existing knowledge of signaling and regulatory networks (29). We applied NicheNet to model interactions between podocytes and GECs, and mesangial cells and GECs that could potentially induce differentially expressed genes (target genes) in GECs in the setting of podocyte injury (Figure 4A). For this analysis, we combined all clusters of each cell type into a single cluster. The top predicted ligands expressed by mesangial cells and/or podocytes were Ptn, Angpt2, Col4a1, Vcam1, Bmp4, Efnb1, Ctgf and Sema3e (Figure 4B). The expression pattern of GEC receptors through which these ligands are known to act were mapped next to the ligand activity analysis. We also mapped the predicted target genes that were differentially expressed in iCTCF^pod-/-^ GECs compared to wildtype controls. The results in Figure 4B are summarized as circular plots in Figures 4C and 4D.

**Figure 4.**
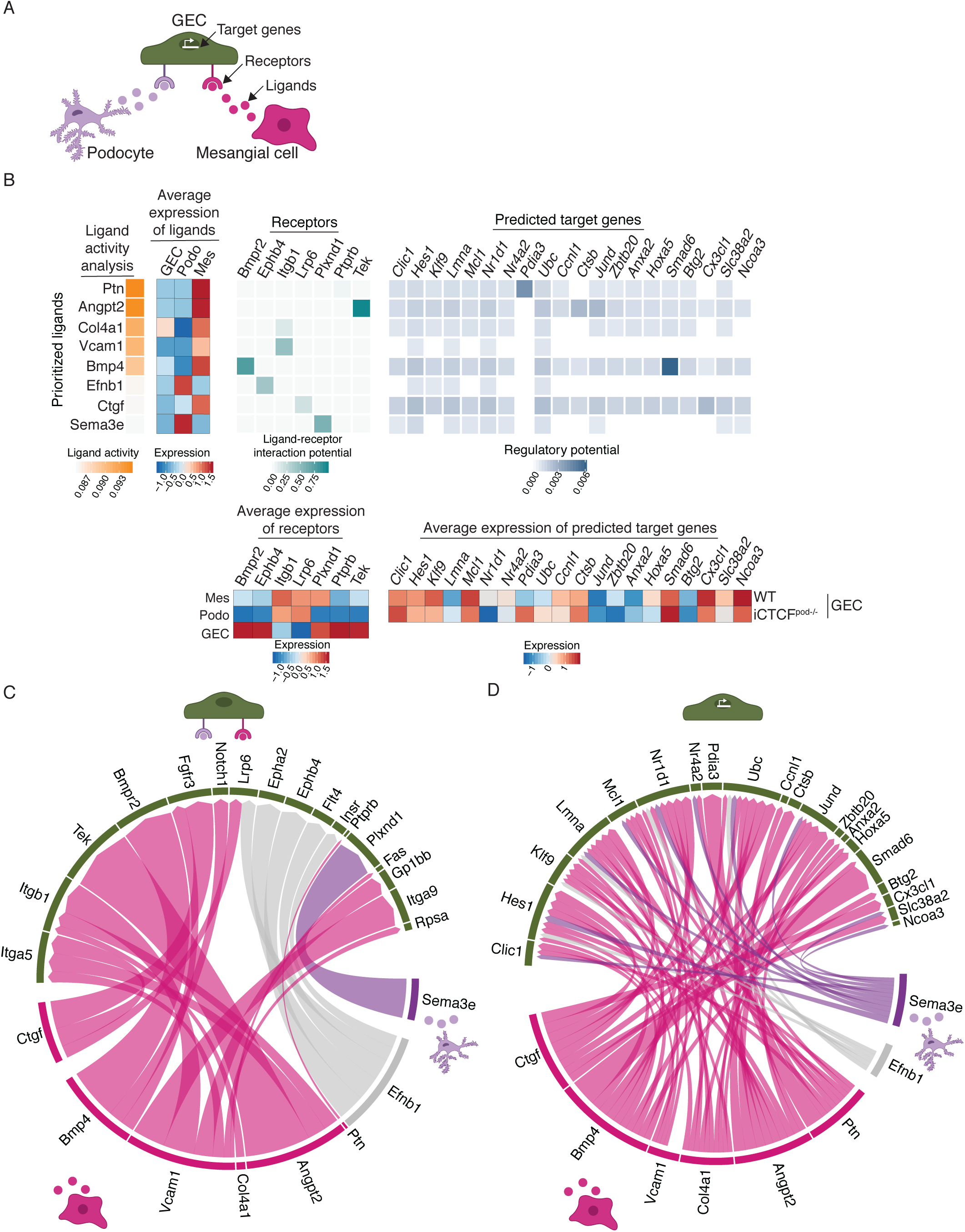
NicheNet analysis reveals ligands, receptors and target genes that contribute to transcriptional changes in GECs as a consequence of podocytes injury. (A) Model for the NicheNet analysis. Potential target genes iCTCF^pod-/-^ GECs were defined as differentially expressed (compared to WT GECs) if they were expressed in at least 10% of cells, had a log fold change of 0.1 and an adjusted p-value of < 0.05. Potential ligand-receptor pairs that lead to differentially expressed genes in iCTCF^pod-/-^ GEC were then identified. (B) Results of the NicheNet analysis. Ligands expressed in mesangial cells and/or podocytes were ranked by the likelihood that the ligand would affect gene expression changes in GECs. Receptors, expressed in GECs, were selected based on their known potential to interact with the prioritized ligands. Finally, target genes were selected based on their differential expression in GECs and their potential to be regulated by the ligand-receptor interactions identified between GECS-podocytes and GECs-mesangial cells. (C) Summary of the ligand-receptor interactions identified in B. (D) Summary of the ligand-target interactions identified in B.

Several ligands, receptors and target genes were notable from this analysis. Pleiotrophin (Ptn), a ligand highly expressed in mesangial cells (Figure 4B), is a secreted growth factor that can bind and inhibit Protein Tyrosine Phosphatase Receptor Type B (Ptprb), stimulating endothelial cell migration via increased Tek (Tie2)/Angpt1 (30) and Kdr (Vegfr2)/Vegfa (31) signaling. Accordingly, Tek was significantly upregulated in GECs (Figure 4B). Angiopoietin-2 (Angpt2), highly expressed in mesangial cells, is an antagonistic ligand of Tek, inhibiting the binding of Angpt1 (32). Since Angpt1 signaling is known to promote podocyte survival and a disruption of the Angpt1/Angpt2 ratio contributes to the development of DKD (33), this ligand-receptor analysis appeared to point to maladaptive Tek signaling in GECs as one of the earliest consequences of podocyte injury.

We also found prominent changes in type IV collagen expression in several glomerular cells (Figure 4B). Normally produced by healthy podocytes, collagen IV heterotrimers transition from □1□1□2 to □3□4□5 during glomerular development, which is necessary for proper GBM formation and function (25, 26). Upregulation of *Col4a1*, along with expansion of the mesangial matrix is frequently observed in DKD (34, 35). In our analysis, *Col4a1* was expressed by both mesangial cells and GECs (average expression of ligand heatmap; Figure 4B), and was specifically upregulated in iCTCF^pod-/-^ GECs compared to wildtype controls. These data suggest that GECs upregulate *Col4a1* in the setting of podocyte injury, which may lead to a stiffer, fibrotic GBM that contributes to the development of segmental sclerosis (25).

Bmp4 was a highly expressed mesangial ligand identified by NicheNet analysis (Figure 4B-D). Bone morphogenetic proteins (BMPs) play key roles in kidney development and disease (36). Bmp4 is upregulated in the setting of diabetic nephropathy in rats (35) and treatment of diabetic mice with an anti-BMP4 antibody prevents the upregulation of *Col4a1* and mesangial matrix expansion (37). Of particular interest, we found that mesangial Bmp4 triggered GEC Smad6 expression (Figure 4B). Expression of Smad6 is induced by BMPs and in a negative-feedback loop, Smad6 specifically inhibits BMPs, including BMP4 (38). Smad6 was downregulated in iCTCF^pod-/-^ GECs compared to wildtype controls, suggesting increased BMP signaling in endothelial cells in the setting of podocyte injury.

One of the prioritized ligands identified in podocytes was Sema3e (Figure 4B-D). Class 3 semaphorins are secreted proteins that function in a variety of biological processes, including angiogenesis, lymphangiogenesis and disease (39). Sema3a is upregulated in human DKD (40) and Sema3g was recently identified as a podocyte-specific gene that protects podocytes from inflammation *in vivo* (41). The role of Sema3e in the podocyte remains unclear. Our analysis revealed a previously unrecognized receptor-ligand pair whereby podocyte Sema3e interacts with plexinD1 on GECs to trigger several downstream gene targets in the setting of podocyte injury (Figure 4C, D).

Among several newly identified GEC target genes, *Hes1* and *Cx3cl1* were also found in the adhesion gene programs enriched in GEC-1 (Figure 4B-D; Supplemental Figure 6). Hes1 is a target of Notch signaling, which plays an important role in the developing kidney, but reactivation can lead to fibrosis (42). Cx3cl1 is a chemokine mainly produced by glomerular endothelium that acts as a chemoattractant and adhesion molecule for its receptor, Cx3cr1, which is ubiquitously expressed on mononuclear and circulatory lymphatic leukocytes (43). Cx3cl1 has been implicated in a variety of kidney diseases, including DKD, IgA nephropathy and glomerulonephritis (43). *Hes1* and *Cx3cl1* were both upregulated in iCTCF^pod-/-^ GECs compared to wildtype controls, suggesting that GECs respond to podocyte injury by upregulating pro-inflammatory and pro-fibrosis programs.

Finally, since many of the ligands and targets identified in the NicheNet analysis are implicated in DKD, in both rodent models as well as humans, we compared our dataset to a single nuclei transcriptomic dataset of early human DKD (10). We specifically compared differentially expressed genes between human and mouse cell clusters. Of the 138 differentially expressed genes in human DKD GECs, 25 genes were also identified in mouse iCTCF^pod-/-^ endothelial cells, including *Col4a1* (Supplemental Figure 7; Supplemental Table 11). Together, these results suggest that endothelial cell programs frequently disrupted in human DKD could be partially recapitulated in our mouse model of early glomerular injury via selective podocyte ablation. In addition, these results suggest that, in addition to podocytes, there may be potential endothelial-specific targets and therapeutic strategies to halt or slow the progression of complex and highly prevalent kidney diseases such as diabetic kidney disease.

## Discussion

Podocyte dysfunction and loss are the cause of many progressive kidney diseases. In this study, we used scRNAseq to dissect the transcriptional networks and cellular crosstalk in the glomerular compartment shortly after induction of selective podocyte injury. We took advantage of a mouse model of inducible podocyte-specific CTCF deletion (iCTCF^pod-/-^) at one week post-induction to generate a single cell map of the kidney filter at the earliest stages of podocyte injury. This comprehensive study of glomerular cells, the first to evaluate the effects of a specific and selective genetic perturbation, has revealed four important insights about the kidney filter in health and disease.

First, we chose to perform scRNAseq after one week of doxycycline treatment to induce CTCF deletion in iCTCF^pod-/-^ animals in order to detect the earliest transcriptional changes that occur after podocyte injury. Although podocyte quantification by image analysis revealed a statistical difference in the number of podocytes in WT and iCTCF^pod-/-^ glomeruli after two weeks of doxycycline treatment (13), here we established that there is a reduction of podocytes in iCTCF^pod-/-^ glomeruli even after one week of doxycycline induction. This suggests that disruption of transcriptional programs critical for podocyte survival precedes histologically detectable podocyte injury and loss, and highlights the power of scRNAseq in discerning subtle changes that can be missed by histology. At a high level, this suggests that efforts to develop podocyte-protective therapies should focus on preventing podocyte injury as early as possible in the course of the disease.

Second, enriching for glomeruli from whole kidneys using the sieving method allowed us to increase the proportion of isolated glomerular cells that were recovered, including a rare population of PECs that have not been captured in previous scRNAseq studies (9, 14, 44). In addition, we were able to identify all major cell types of the kidney. This provides the opportunity for future studies to evaluate transcriptional changes driven by podocyte injury in the entire kidney, which will deepen our understanding of kidney-wide transcriptional events leading to the progression of CKD.

Third, our data suggests that the iCTCF^pod-/-^ mouse model recapitulates many aspects of human disease at the transcriptional level, including differential expression of genes critical for podocyte adherence to the GBM and autocrine pro-survival ligands. Since adherence of podocytes to the GBM is critical for their function, the upregulation of *Col4a5* and other genes in cell adhesion gene programs support a model whereby podocytes respond to initial injury by increasing adherence to the GBM. Human mutations in podocyte focal adhesion genes, including actin cytoskeleton genes, type IV collagen and integrins have all been implicated in FSGS and/or nephrotic syndrome (45, 46). Further, ligand-receptor analysis revealed that signaling molecules implicated in human DKD are also disrupted in the CTCF mouse model (eg. BMP4, Col4a1) (34, 35, 37). We also observed the dysregulation of two key autocrine pro-survival ligands, *Vegfa* and *Pdgfb* whose roles have been well established *in vitro* and *in vivo*, and alterations in their expression and/or activity have been associated with human kidney diseases such as thrombotic microangiopathies and DKD (47). Specifically, anti-VEGF therapy, a common cancer treatment, leads to renal thrombotic microangiopathy (48), and *Pdgfb* expression is increased in human kidney biopsies of patients with DKD and mesangial proliferative glomerulonephritis (49–51). Taken together, these data support the notion that the iCTCF^pod-/-^ mouse model may offer a reasonably faithful transcriptional profile of early glomerular changes with relevance to several human kidney diseases.

Our detailed modelling of ligand-receptor-target gene interactions in all glomerular cells after podocyte injury led to several key observations with important therapeutic implications. We found that GECs were most affected by podocyte injury, consistent with the physical proximity between the two cell types. This result emphasizes the importance of moving beyond our reductionist studies focusing on one cell type alone, by understanding intercellular communication patterns in order to gain deeper insights into mechanisms of kidney disease progression. Our integrated analysis revealed several previously unrecognized molecules in GECs, such as BMP4 and PlexinD1, and in podocytes, such as Sema3E, that may be further explored for their potential to be targeted for therapeutic benefit. In this context, the detailed single cell map of the mouse glomerulus after early podocyte injury generated in this study, made openly available to all investigators, may be used to generate hypotheses about actionable targets for much needed therapies.

## Methods

### Animal care

All animal studies were performed in accordance with guidelines established and approved by the Animal Care and Use Committee at Brigham and Women’s Hospital, Harvard Medical School. iCre^pod^-Ctcf^wt/fl^ mice were generated as previously described (13) and inbred to generate iCre^pod^-Ctcf^fl/fl^ (iCTCF^fl/fl^) and iCre^pod^-Ctcf^wt/wt^ (WT) mice. Both male and female mice were used in this study. Doxycycline (4 g/L) (Sigma D9891) was continuously administered in drinking water containing sucrose (50 g/L) (VWR BDH0308) to both iCTCF^fl/fl^ (generates iCTCF^pod-/-^ mice) and WT littermate control mice (age 6-8 weeks at start of doxycycline) to drive Cre expression specifically in podocytes.

### Glomerular enrichment from whole kidney

iCTCF^pod-/-^ and WT mice were sacrificed after 1 week of doxycycline treatment and kidneys were quickly dissected and washed with ice-cold Hanks’ Balanced Salt Solution (HBSS; Thermo Scientific 14170112). After removing the kidney capsules, glomeruli were enriched at 4 °C using the sieving technique (52) with 180 ⎧m, 75 ⎧m and 53 ⎧m sieves. Glomerular-enriched fractions collected from the 53 ⎧m sieve were rinsed with ice-cold 1X HBSS and placed on ice.

### Preparation of single-cell suspensions

Glomerular-enriched fractions were centrifuged at 350 xg for 5 mins at room temperature. After removing most of the HBSS, 1 ml liberase TH digestion buffer (Sigma 5401135001) containing 50 U/⎧l DNase I (Thermo Scientific 90083) was added to the glomerular pellet and incubated at 37 °C for 60 minutes on an orbital shaker (500 rpm). The suspension was passed through a 27-gauge needle 2x after 20 minutes. Digested glomerular fractions were added to 9 ml RPMI (Thermo Scientific 11875119) containing 10% FBS (Thermo Scientific 16000044) and centrifuged at 500 xg for 5 minutes at room temperature. 1 ml red blood cell lysis buffer (Thermo Scientific A1049201) was added to the glomerular pellet and mixed for 1 minute at room temperature. 9 ml RPMI (10% FBS) was added and the suspension was centrifuged at 500xg for 5 minutes at room temperature. The media was removed from the pellet and 200 ⎧l Accumax (STEMCELL technologies 07921) was added and incubated for 20 mins at 37 °C. 1.8 ml 1X PBS (Thermo Scientific 14190250) + 0.04% BSA (Sigma A1933) was added and the suspension was centrifuged at 500 xg for 8 minutes. The digested glomeruli were washed with 750 ⎧l 1X PBS + 0.04% BSA and filtered using a 40 ⎧m Flowmi Tip Strainer (Sigma BAH136800040). 1.25 ml 1X PBS + 0.04% BSA was added and the suspension was centrifuged at 500 xg for 8 minutes. The pellet was resuspended in a small volume of 1X PBS + 0.04% BSA.

### Library preparation and single-cell sequencing

Single cells were processed through the 10X Chromium 3’ Single Cell Platform using the Chromium Single Cell 3’ Library, Gel Bead and Chip Kits (10X Genomics, Pleasanton, CA), following the manufacturer’s protocol. Briefly, 10,000 cells were added to each channel of a chip to be partitioned into Gel Beads in Emulsion in the Chromium instrument, followed by cell lysis and barcoded reverse transcription of RNA in the droplets. Breaking of the emulsion was followed by amplification, fragmentation and addition of adapter and sample index. Libraries were pooled together and sequenced on Illumina HiSeq.

### Hybridization Chain Reaction (HCR)

All HCR v3 reagents (probes, hairpins, and buffers) were purchased from Molecular Technologies. Thin sections of tissue (10 ⎧m) were mounted in 24-well glass bottom plates (VWR 82050-898) coated with a 1:50 dilution of APTES (Sigma 440140). The following solutions were added to the tissue: 10% formalin (100503-120, VWR) for 15 minutes, 2 washes of 1X PBS (ThermoFisher Scientific AM9625), ice cold 70% EtOH at - 20°C for 2 hours (to overnight), 3 washes of 5X SSCT (ThermoFisher Scientific 15557044, with 0.2% Tween-20), Hybridization buffer (Molecular Technologies) for 10 minutes, probes in Hybridization buffer overnight, 4 washes of Wash buffer (Molecular Technologies) for 15 minutes, 3 washes of 5X SSCT, Amplification buffer (Molecular Technologies) for 10 minutes, hairpins were heat denatured in Amplification buffer overnight, 3 washes of 15 minutes with 5X SSCT (1:10,000 DAPI TCA2412-5MG, VWR in the second wash), and storage/imaging in 5X SSCT. Imaging was performed on a spinning disk confocal (Yokogawa W1 on Nikon Eclipse Ti) operating NIS-elements AR software. Image analysis and processing was performed on ImageJ Fiji.

## Statistics

### Data processing and quality control

We used the Cellranger toolkit (v2.1.1) to perform de-multiplexing using the “cellranger mkfastq” command, and the “cellranger count” command for alignment to the mouse transcriptome, cell barcode partitioning, collapsing unique molecular identifier (UMI) to transcripts, and gene-level quantification. We filtered cells to include cells expressing a minimum of 500 genes and a maximum of 4000 genes. Further, the percentage of reads mapping to mitochondrial genes was capped at 12%. We used DoubletFinder to identify potential doublets (53). Clusters with over 75% of cells classified as “high-confidence” doublets were removed from further analysis. The remaining cells classified as “high-confidence” doublets were also removed from further analysis.

### Unsupervised clustering and dimensionality reduction

The default settings in the Seurat R package (54) (v3.0) were used for normalization (NormalizeData) of the gene expression counts, identifying variable genes (FindVariableGenes), finding integration anchors (FindIntegrationAnchors) and integrating the samples (IntegrateData). Unwanted variation due to number of UMIs and ratio of reads mapping to mitochondrial genes (ScaleData). Dimensionality reduction was performed using Principal Component Analyses (RunPCA) on the highly variable genes. To distinguish principal components (PCs) for further analysis, we used the PCElbowPlot() function. For clustering all cells, the first 40 PCs sufficiently captured all of the variance. We identified molecularly distinct clusters using the default parameters (FindClusters) and a resolution of 0.4 (FindNeighbors). After subsetting glomerular cells, we processed and scaled the data as described above. We used 30 PCs for downstream clustering and a resolution of 0.3 (FindNeighbors).

### Cell type classification

Cluster-enriched or marker genes were computed using the Wilcoxon-Rank sum test (FindAllMarkers) for differential expression of genes in the cluster cells vs all other cells and selecting those genes that pass the adjusted p-value (FDR) cutoff of 0.05 as cluster-representative. Cluster identity was assigned by comparing data-driven genes with a list of literature-curated genes for mature kidney cell types (Supplemental Table 3).

### Differential gene expression analysis

To analyze differential expression between iCTCF^pod-/-^ and WT cells in a specific cluster, we performed pair-wise differential expression analysis in Seurat (FindMarkers) with logFC.threshold set to 0.01 and default parameters. Within “FindMarkers”, “ident.1” was set to cluster-specific iCTCF^pod-/-^ cells and “ident.2” was set to the corresponding cluster-specific WT cells. Genes with an adjusted p-value < 0.05 were considered significant.

### Over-representation analysis

Over-representation analysis was used to determine whether any known biological functions or processes were over-represented, or enriched, in our list(s) of differentially expressed genes (55). We used the R package clusterProfiler (56), specifically the enrichGO function, for the over-representation analysis. Default parameters were used, with the exceptions of ont=“BP”, pvaluecutoff = 0.25 and qvaluecutoff = 0.25. We then used the R package enrichplot (https://github.com/YuLab-SMU/enrichplot), specifically the emapplot function, to visualize the enrichment results by plotting the top enrichment terms.

### Ligand receptor analysis

iTALK (57) was used to map receptors to ligands that were differentially expressed in iCTCF^pod-/-^ podocytes. Specifically, “rawParse” was used to calculate the mean expression of each gene, using scaled data from Seurat. “FindLR” using datatype=“meancount” was used to identify ligand/receptor pairs. Ligand/receptor pairs of interest were plotted using default parameters of “LRPlot”. The thickness of the lines indicates the relative mean expression of the ligand and the size of the arrowhead indicates the relative mean expression of the receptor.

### NicheNet Analysis

We used the R package NicheNet (29) to predict ligand-receptor interactions that might drive gene expression changes in our cell type of interest. We combined all podocyte clusters and all endothelial clusters for this analysis. All default parameters were used with the exception of setting a lower cutoff threshold of 0.11 for “prepare_ligand_target_visualization”.

### HCR image processing and quantification

Images were processed using ImageJ2 (National Institutes of Health). Raw ND2 files were background subtracted using the Rolling Ball method (rolling=50 sliding stack). Average intensities of the Z-stack images were then projected, and image channels were split and saved separately. CellProfiler (version 3.1.5, Broad Institute) (58) was used for cell segmentation based on the fluorescence intensity of DAPI channel and for measuring integrated fluorescence intensity the rest of the channels. Four WT and four iCTCF^pod-/-^ mice were evaluated for statistical analysis. All glomeruli from three 40X images were quantified. A Welch-corrected two-tailed t-test was performed.

## Study approval

This study was approved by the Animal Care and Use Committee at Brigham and Women’s Hospital, Harvard Medical School.

## Supporting information

Supplemental Table 1-4: Quality control metrics of single-cell RNA sequencing samples

Supplemental Table 5: Percentage of podocytes

Supplemental Table 6: Log fold changes

Supplemental Table 7: Differentially expressed genes

Supplemental Table 8: Over-representation analysis

Supplemental Table 9: Uniquely differentially expressed genes

Supplemental Table 10: Uniquely differentially expressed genes in GEC-1

Supplemental Table 11: Differentially expressed genes in human DKD GECs

## Author Contributions

AC, JM and AG designed the experiments; AC performed all animal experiments, assisted by YZ and MM. AC performed all computational analysis of single-cell RNA sequencing data; LN generated and sequenced 10X libraries; JM performed all HCR experiments and HC performed HCR computational analysis for quantification. AG directed the project. AC and AG wrote the manuscript. All authors read, commented and approved the manuscript.

## Acknowledgements

We are grateful to Aviv Regev and Orit Rosenblatt-Rosen and members of the Regev group for ongoing advice on single cell technology and computational approaches. We thank the Broad GP platform for sequencing services.

## Conflict of Interest

None relevant to this work

## Figure legends

**Supplemental Figure 1.**
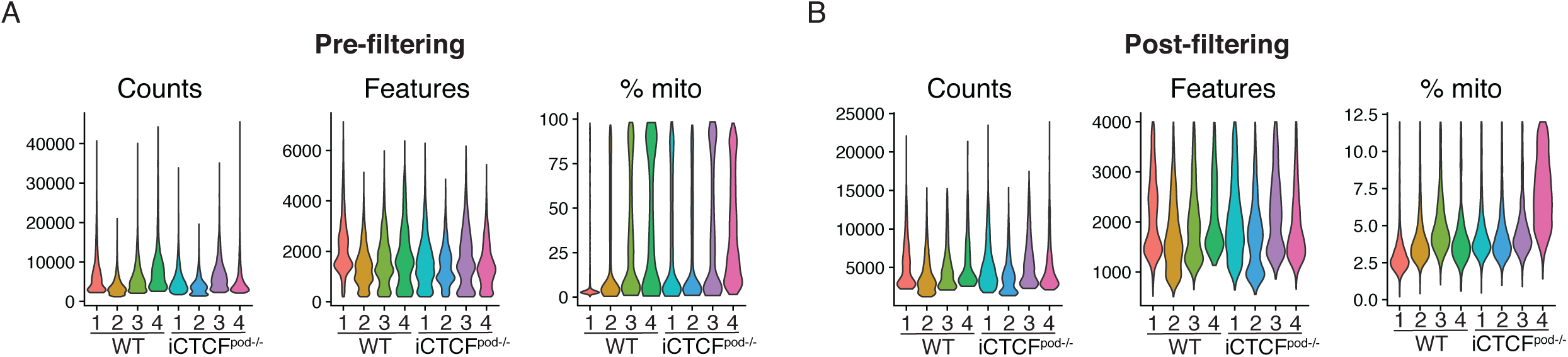
Quality control metrics of single-cell RNA sequencing data. (A,B) Violin plots showing the distribution of the number of unique molecular identifiers (Counts), number of genes (Features) and percentage of mitochondrial reads (% mito) in each sample before (A) and after (B) filtering.

**Supplemental Figure 2.**
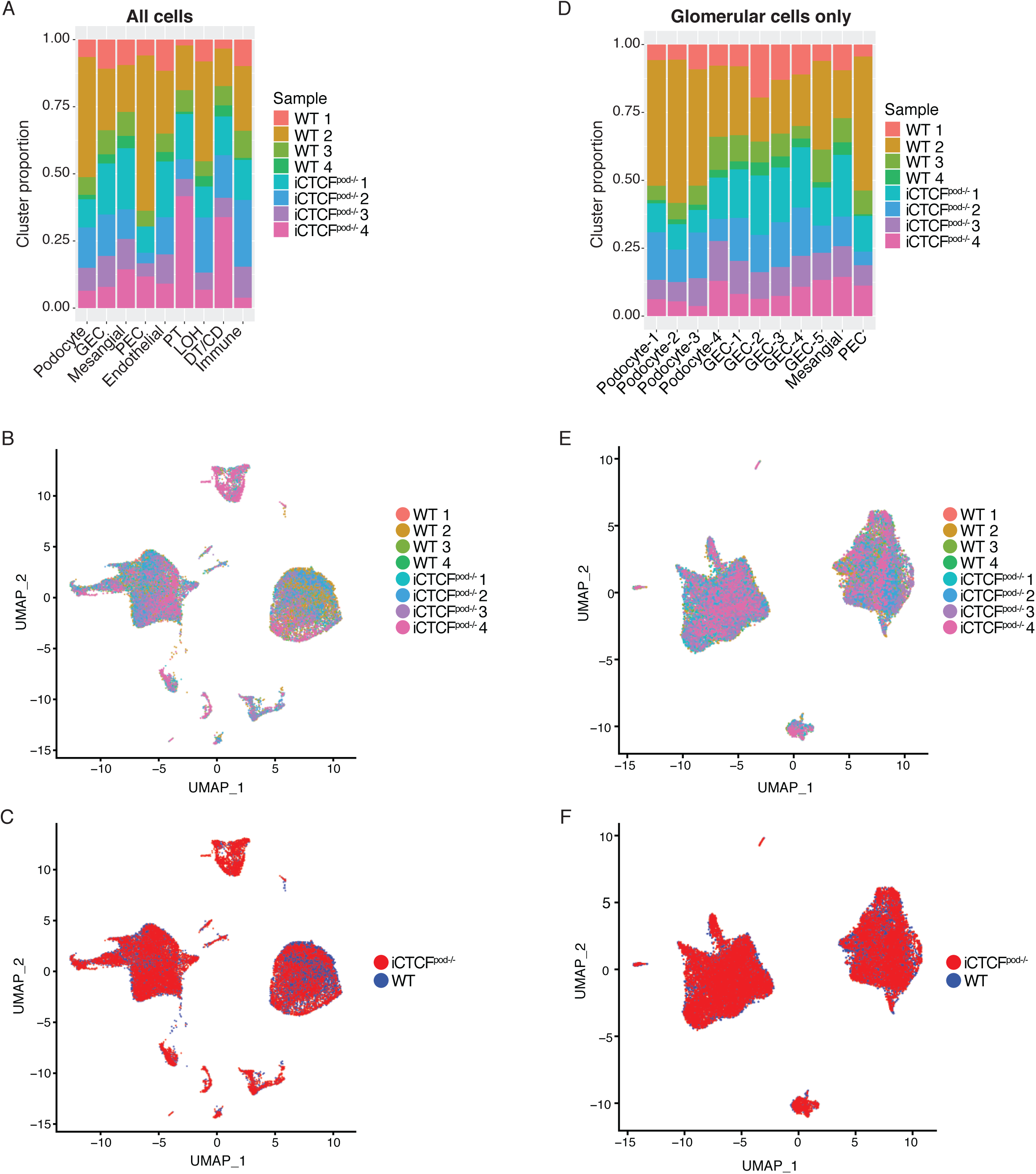
Clustering of WT and iCTCF^pod-/-^ cells by sample and genotype. (A-C) All profiled WT and iCTCF^pod-/-^ cells. (A) Distribution of all profiled WT and iCTCF^pod-/-^ cells per cluster. (B-C) Clustering of all WT and iCTCF^pod-/-^ cells colored by sample number (B) and genotype (C). (D-F) Isolated and reclustered WT and iCTCF^pod-/-^ glomerular cells. (D) Distribution of WT and iCTCF^pod-/-^ glomerular cells per cluster. (E-F) Clustering of WT and iCTCF^pod-/-^ glomerular cells colored by sample number (E) and genotype (F).

**Supplemental Figure 3.**
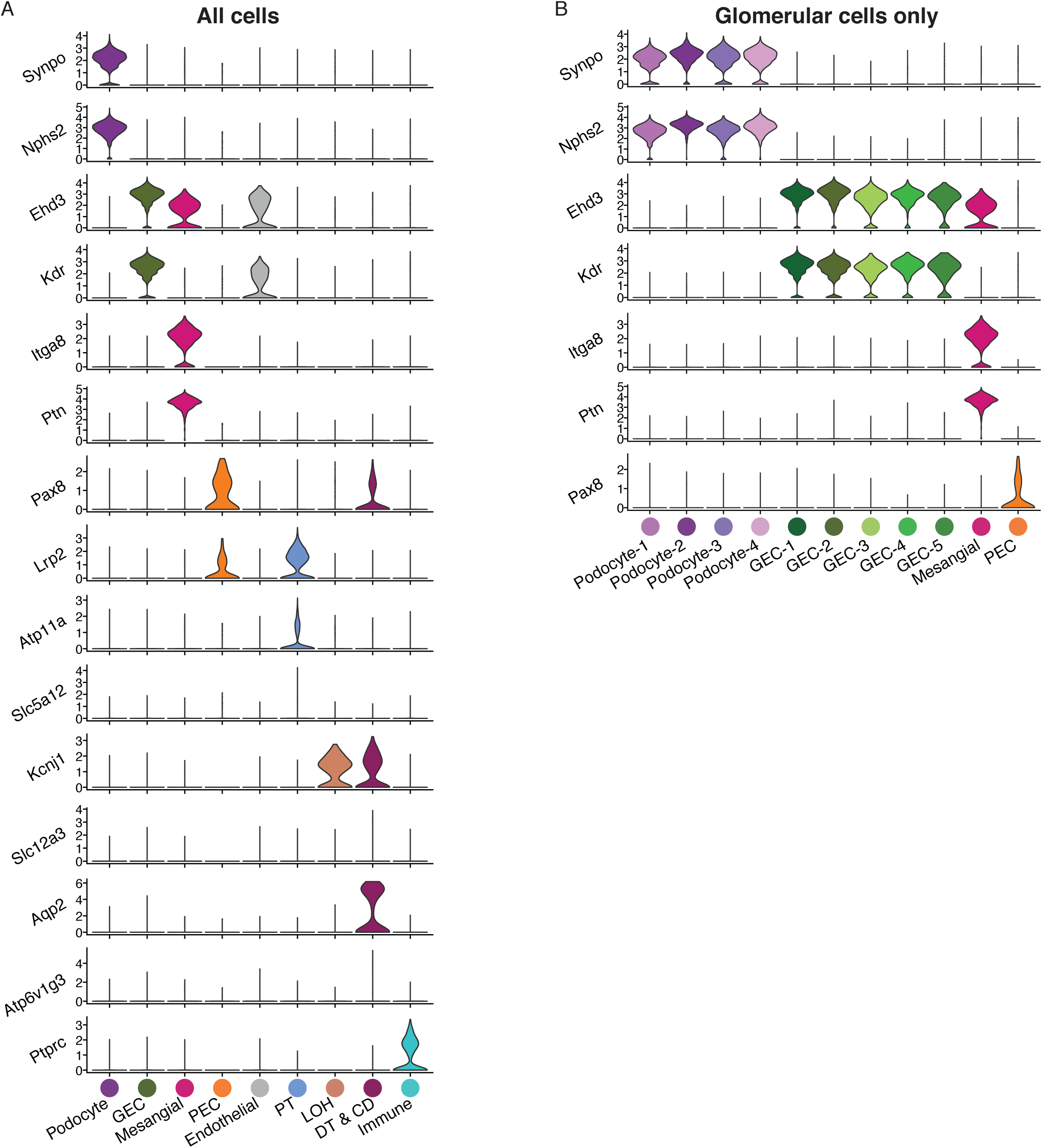
Expression of cell-type specific markers in WT and iCTCF^pod-/-^ cells used for cluster identification in Figure 1. Violin plots of cell-type specific markers in WT and iCTCF^pod-/-^ cells used for cluster identification for (A) all cells or (B) glomerular cells only.

**Supplemental Figure 4.**
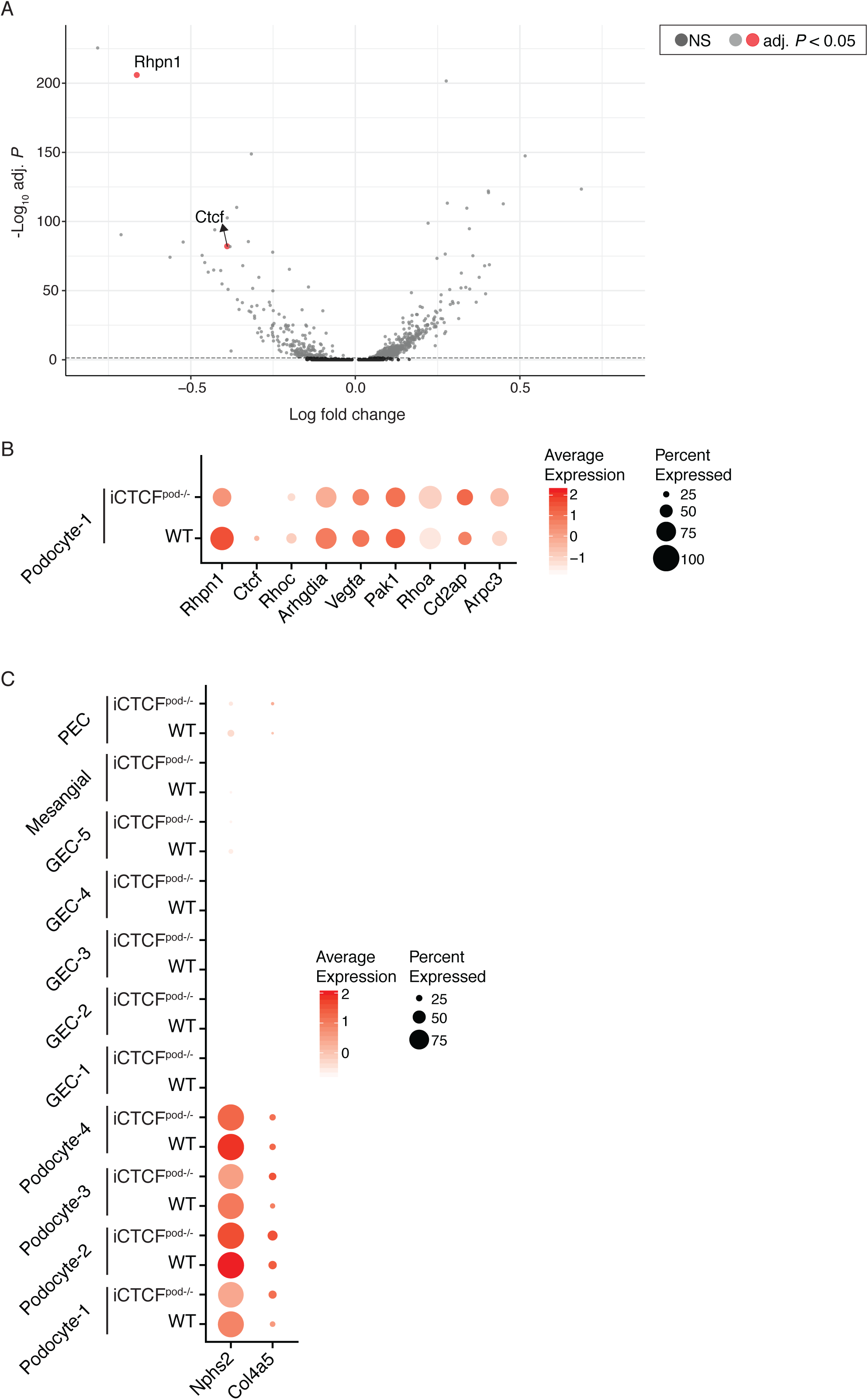
Extension of Figure 2. (A) Full volcano plot presented in Figure 2, highlighting *Rhpn1* and *Ctcf*. Significantly differentially expressed genes are represented by light grey and red dots. Genes in dark grey are not significantly differentially expressed. (B) Dot Plot showing the expression of all labeled genes in Volcano plots of podocyte-1 cells. Size of the dot represents the percent of cells expressing the genes. Color represents the average expression level of the gene in the specified cluster. (C) Dot Plot showing the expression of *Nphs2* and *Col4a5* in all clusters.

**Supplemental Figure 5.**
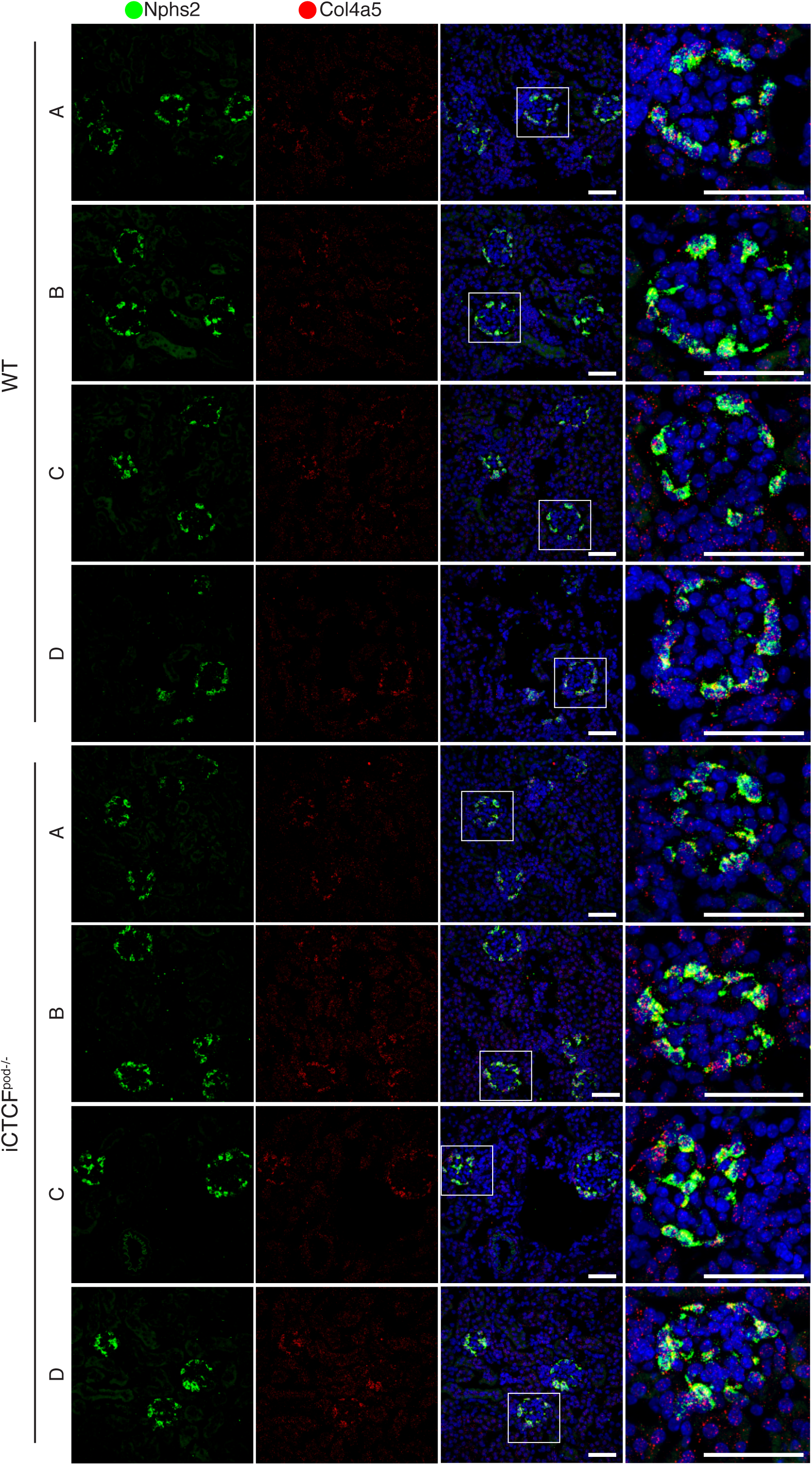
Extended validation of genes presented in Figure 3. Additional HCR images used for quantification of *Nphs2* and *Col4a5* expression in WT and iCTCF^pod-/-^ animals. Scale bars are 50 μm.

**Supplemental Figure 6.**
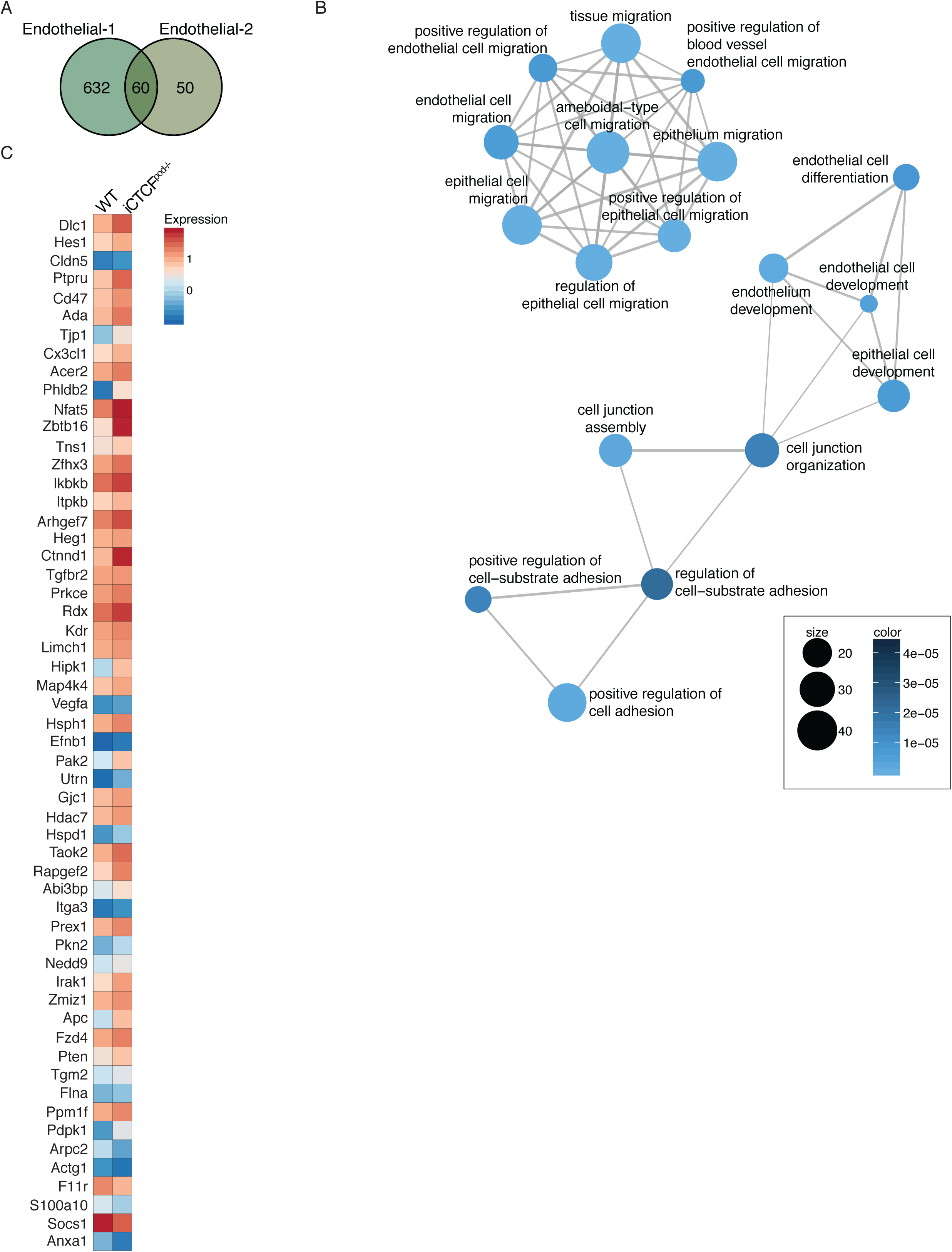
Endothelial cells enriched for gene programs important for adhesion and migration. (A) Venn diagram of the differentially expressed genes identified in GEC-1 and GEC-2. (B) Visualization of the over-representation analysis of genes uniquely differentially expressed in GEC-1 presented as a network, or enrichment map. Overlapping gene sets cluster together. (C) Heatmap of all genes from the adhesion and migration gene programs from B.

**Supplemental Figure 7.**
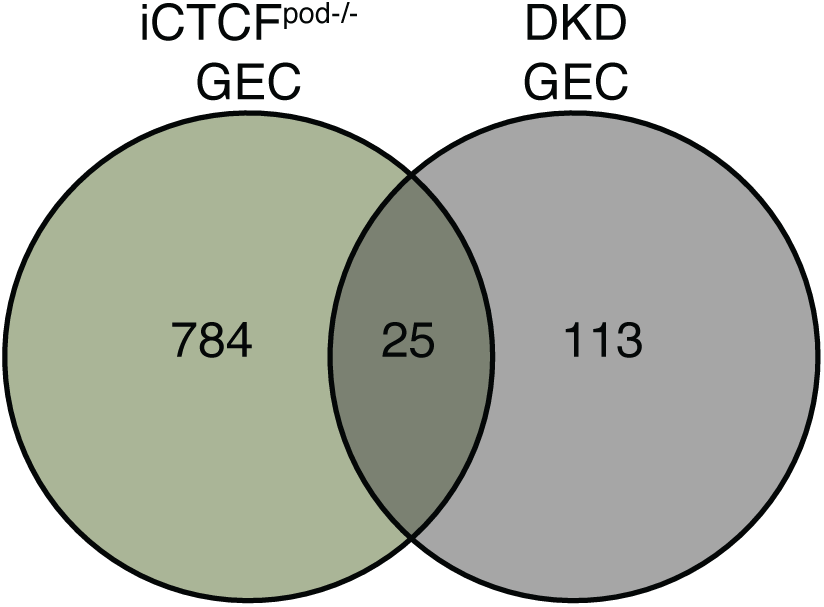
Comparison of iCTCF^pod-/-^ GECs with human DKD GECs. Venn diagram of the differentially expressed genes found in all iCTCF^pod-/-^ GEC clusters and human DKD GEC clusters.

## Notes

### Competing Interest Statement

The authors have declared no competing interest.

