## Supplemental Table 1-4: Quality control metrics of single-cell RNA sequencing samples for "Single cell transcriptomics reveal disrupted kidney filter cell-cell interactions after early and selective podocyte injury"

Supplemental Table 1. Quality control metrics of single-cell RNA sequencing samples.

| Sample | # Reads | Median Genes  per Cell | Sequencing  Saturation | Estimated  # Cells | Fraction Reads  in Cells |
| --- | --- | --- | --- | --- | --- |
| WT-1A | 189,112,059 | 1,993 | 85.7% | 1,917 | 80.0% |
| WT-1B | 207,773,772 | 1,818 | 88.2% | 1,831 | 79.3% |
| WT-2A | 136,379,215 | 1,016 | 61.4% | 9,893 | 83.2% |
| WT-2B | 147,236,638 | 1,185 | 59.9% | 8,967 | 84.1% |
| WT-3 | 218,252,929 | 1,293 | 66.4% | 7,132 | 77.6% |
| WT-4A | 257,025,194 | 1,438 | 92.3% | 1,202 | 70.1% |
| WT-4B | 250,252,386 | 1,463 | 92.0% | 1,196 | 69.0% |
| iCTCF^pod-/-^ -1A | 180,179,138 | 1,172 | 69.3% | 6,424 | 79.4% |
| iCTCF^pod-/-^ -1B | 197,042,703 | 1,287 | 71.9% | 5,748 | 77.4% |
| iCTCF^pod-/-^ -2 | 151,216,812 | 1,144 | 62.8% | 8,782 | 82.2% |
| iCTCF^pod-/-^ -3A | 231,503,612 | 1,315 | 75.8% | 4,783 | 76.3% |
| iCTCF^pod-/-^ -3B | 218,073,195 | 1,347 | 74.0% | 4,760 | 76.2% |
| iCTCF^pod-/-^ -4A | 179,510,830 | 1,179 | 50.8% | 7,959 | 57.7% |
| iCTCF^pod-/-^ -4B | 189,898,909 | 1,257 | 54.3% | 7,076 | 56.3% |

Supplemental Table 2. Cell number per sample before and after each filtering step.

| **Filtering Step** | **WT cells remaining** | | | | **iCTCF^pod-/-^ cells remaining** | | | |
| --- | --- | --- | --- | --- | --- | --- | --- | --- |
|  | 1 | 2 | 3 | 4 | 1 | 2 | 3 | 4 |
| Original | 3688 | 15495 | 6215 | 2132 | 10316 | 7730 | 8079 | 13259 |
| Low and high gene count | 3276 | 13623 | 5189 | 1619 | 8659 | 6937 | 6377 | 11261 |
| High mitochondrial content | 2659 | 9846 | 2489 | 779 | 4887 | 4659 | 3103 | 3086 |
| DoubletFinder | 2500 | 9202 | 2346 | 735 | 4551 | 4364 | 2924 | 2888 |

Supplemental Table 3. Markers used for cluster identification.

| Gene | Cell Type |
| --- | --- |
| Synpo | Podocyte |
| Nphs2 | Podocyte |
| Ehd3 | GEC |
| Kdr | GEC |
| Itga8 | Mesangial |
| Ptn | Mesangial |
| Pax8 | PEC |
| Lrp2 | PT |
| Atp11a | PT-S3 |
| Slc5a12 | PT-S1 |
| Kcnj1 | LOH |
| Slc12a3 | DT |
| Aqp2 | CD-PC |
| Atp6v1g3 | CD-IC |
| Ptprc | Immune |

Supplemental Table 4. Number of cells per genotype and per cluster in Figure 1C.

| Cell type | Genotype | # cells |
| --- | --- | --- |
| Podocyte-1 | WT | 3011 |
| Podocyte-1 | iCTCF^pod-/-^ | 2138 |
| Podocyte-2 | WT | 1538 |
| Podocyte-2 | iCTCF^pod-/-^ | 788 |
| Podocyte-3 | WT | 1115 |
| Podocyte-3 | iCTCF^pod-/-^ | 716 |
| Podocyte-4 | WT | 607 |
| Podocyte-4 | iCTCF^pod-/-^ | 633 |
| GEC-1 | WT | 3679 |
| GEC-1 | iCTCF^pod-/-^ | 4327 |
| GEC-2 | WT | 1233 |
| GEC-2 | iCTCF^pod-/-^ | 1325 |
| GEC-3 | WT | 370 |
| GEC-3 | iCTCF^pod-/-^ | 448 |
| GEC-4 | WT | 189 |
| GEC-4 | iCTCF^pod-/-^ | 311 |
| GEC-5 | WT | 79 |
| GEC-5 | iCTCF^pod-/-^ | 71 |
| Mesangial | WT | 309 |
| Mesangial | iCTCF^pod-/-^ | 452 |
| PEC | WT | 101 |
| PEC | iCTCF^pod-/-^ | 59 |
